## Supplementary material for "BRASS: permutation methods for binary traits in genetic association studies with structured samples": S1 Text

### Simulation studies: Genotypes

The Balding-Nichols model [1] is used to draw allele frequencies for 2 sub-populations with the fixation index $F$ set to 0.01 which is representative of the population structure seen in humans within continents [2]. For each marker $s$, given the ancestral allele frequency $p_s$ drawn from a uniform distribution on $[0.2, 0.8]$ (and independently across markers), the allele frequency in sub-population $k{=}1, 2$ is drawn independently from a Beta distribution with parameters $p_s(1 - F)/F$ and $(1 - p_s)(1 - F)/F$. An equal number of pedigrees (each with structure given in S1 Fig) is assigned to the 2 sub-populations with all founders within a given pedigree being assigned to the same sub-population. For each marker, genotypes for a pedigree's founders are simulated as independent Bernoulli draws using the corresponding sub-population allele frequency, and gene dropping is used to determine genotypes for other pedigree members.

We simulate $M_c = 10,000$ causal SNPs which are used to generate polygenic effects in the trait generative model as well as to correct for correlation in the sample when simulating trait replicates from the seven methods. The $M_c$ causal markers are simulated only once, and they are used to compute the GRM estimate $\tilde{\boldsymbol{\Phi}}$ in (4), which is then reused in all simulations. In addition, independent non-causal markers are simulated for each simulation replicate using the same Balding-Nichols model, where the ancestral allele frequencies are drawn from a uniform distribution on $[0.2, 0.8]$, and these are the markers tested for association.

### Simulation studies: Trait and Covariates

We simulate 46 pedigrees of the same configuration (S1 Fig), each consisting of 22 individuals. For each individual, three covariates are simulated: sex, age and an i.i.d. standard normal covariate. Sex is determined by the pedigree configuration and age is drawn uniformly and independently within 1.5 years of $73, 75, 46, 43, 40, 46, 40, 43, 47, 51, 18, 21, 15, 15, 12, 9, 13, 17, 24, 21, 18$, and 14 years respectively for individuals labeled $1-22$ in the pedigree. All the covariates (besides sex) are re-generated for each simulation replicate.

We consider two generative models for the trait. First, a logistic model is used in which $Y_i$ is given by,

$$Y_i|\mathbf{X}_i, \mathbf{W}_i, \boldsymbol{\alpha} \sim \text{Bernoulli}(p_i), \text{ independently, with } \text{logit}(p_i) = \mathbf{X}_i\boldsymbol{\beta} + \mathbf{W}_i\boldsymbol{\alpha}, \qquad (1)$$

where $\mathbf{X}_i$ is the covariate row vector for the $i$th individual; $\boldsymbol{\beta}$ are the fixed effects for the covariates; $\mathbf{W}_i$ is a row vector representing the SNP information for the $i$th individual corresponding to the $M_c$ causal markers standardized to have mean 0 and variance 1; $\boldsymbol{\alpha} = (\alpha_1, \cdots, \alpha_{M_c})^T \sim MVN(0, \sigma^2\mathbf{I})$ represent the random effects of the $M_c$ causal markers with polygenic variance $\sigma_a^2 = \sigma^2 M_c$. The values for $\boldsymbol{\beta}$ and $\sigma_a^2$ are chosen such that: (1a) If all covariates are included in the fitted model, they each explain an equal amount of the variation on the logit scale due to covariates; (1b) If the standard normal covariate is excluded, its effect size is set to correspond to an average p-value of 0.05 in a LMM Wald test (determine by simulations), and age and sex equally explain the remaining portion of the variability due to covariates; (2) Considering the total variance on the logit scale due to covariate effects and polygenic random effects, the fraction of this variance due to covariates is fixed at 20%, 40%, 60% or 80%; (3) either (a) Bernoulli error explains on average about 20% of the total phenotypic variability and the prevalence is approximately 30% or (b) Bernoulli error explains on average about 55% of the total phenotypic variability and the prevalence is approximately 5%. When ascertainment is included in the sample, we simulate data under the model in (1) and select either (a) 500 cases and 500 controls or (b) 300 cases and 700 controls at random to be retained in the sample.

We also consider generating the trait using a liability threshold model,

$$Y_i = \mathbb{1}_{\{L_i > 0\}}, \text{ with } L_i = \mathbf{X}_i\boldsymbol{\beta} + \mathbf{W}_i\boldsymbol{\alpha} + \epsilon_i, \tag{2}$$

where $L_i$ is the latent liability for individual $i$. The values of $\boldsymbol{\beta}$ and $\sigma_a^2$ are chosen using almost the same conditions as for the logistic model, where conditions (1a), (1b), and (2) are enforced on the liability scale instead of the logit scale, the random error $\epsilon_i$ in Eq (2) explains 20% of the phenotypic variability in all cases, and the prevalence is set to be either (a) 30% or (b) 5%. In both Eq (1) and (2), larger values of $\sigma_a^2$ correspond to more severe confounding effects of population/family structure on phenotype-genotype association. Ascertainment is as described above.

Owing to the computational intensive nature of the simulations, the trait vector is simulated 200 times, and 150 independent marker panels, each consisting of $m = 100$ non-causal markers, are simulated for each trait replicate. Retaining only the top association signal in each panel, this results in 150 association test statistics which overall amounts to 20,000 replicates used for type 1 error estimation.

### Incorporating the sample structure in the null model

In order to get trait replicates, all seven resampling methods involve fitting a prospective model under the null that requires an estimate for the GRM $\boldsymbol{\Phi}$ to capture the genetic relatedness in the sample. Our approach is to model genetic relatedness as both fixed and random effects in the null model, where the fixed component is represented by the inclusion of covariates that represent major axes of genetic variation, and the random component is represented by the inclusion of a GRM that reflects the leftover structure. We use a previously proposed method, PC-Relate [3], where we first estimate the GRM $\boldsymbol{\Phi}$ using population estimates for the minor allele frequencies (MAF), and whose entries are,

$$\widehat{\boldsymbol{\Phi}}_{ij} = \frac{1}{L}\sum_{l=1}^{L}\frac{(G_{il} - 2\hat{p}_l)(G_{jl} - 2\hat{p}_l)}{2\hat{p}_l(1-\hat{p}_l)}, \tag{3}$$

where $L$ is the number of markers, $G_{il}$ is the minor allele count for the $i$-th individual at marker $l$, and $\hat{p}_l = 0.5 \cdot \overline{G}_l$, where $\overline{G}_l$ is the sample average estimator of the population MAF for marker $l$. We obtain the top $D$ PCs from $\widehat{\boldsymbol{\Phi}}$, and these PCs are then included as ancestry informative covariates in the null model. We then build a GRM estimate $\tilde{\boldsymbol{\Phi}}$ to capture the remaining genetic similarities among subjects that are not reflected by the top PCs. The entries of $\tilde{\boldsymbol{\Phi}}$ are,

$$\tilde{\boldsymbol{\Phi}}_{ij} = \frac{\sum\limits_{l=1}^{L}(G_{il} - 2\widetilde{p}_{il})(G_{jl} - 2\widetilde{p}_{jl})}{2\sum\limits_{l=1}^{L}\left[\widetilde{p}_{il}(1-\widetilde{p}_{il})\widetilde{p}_{jl}(1-\widetilde{p}_{jl})\right]^{1/2}}, \tag{4}$$

where $G_{il}$ and $\widetilde{p}_{il}$ are the minor allele count and a predicted subject-specific MAF, respectively, for the $i$-th individual at marker $l$. For each marker $l$, the subject-specific MAF estimate is obtained as half of the fitted values from a linear regression of $\mathbf{G}_l$ on the top $D$ PCs. So in the numerator of (4), the centered genotype values represent the residuals from this linear regression, effectively removing the effects captured through the top $D$ PCs. The resulting $\tilde{\boldsymbol{\Phi}}$ is used as the GRM estimate in all seven resampling methods.

To assess significance in each data simulation, up to 10,000 trait replicates are generated under the null using each of the seven methods. For each marker panel, the trait replicates are individually tested for association with the non-causal markers, and the smallest p-value for each replicate is compared with the one from the original data simulation. We use CARAT [4] for single-SNP association testing; it is a retrospective association test for binary traits in structured samples.

**Adaptive resampling procedure.** Given the computationally intensive nature of permutation-based approaches, we use an adaptive procedure when simulating replicates for multiple testing correction at level $\alpha$, where our chosen stopping criteria are checked initially with 1000 simulated replicates and then in increments of 5000 starting at 5000 replicates. More specifically, we continue generating replicates until either (1) $N_{max}$, the maximum number of replicates allowed, has been reached (where $N_{max} \geq 1000$); (2) A hypothesis test for $H_0 : p = \alpha$ against $H_a : p > \alpha$ is rejected at significance level 0.01, where the estimate for the p-value $p$ is the proportion of replicates with test statistic at least as extreme as the one observed. An exact test for $H_0$ is used when the number of replicates is 1000; otherwise, a z-test is performed. If $N_{max}$ has been reached, the p-value estimated from the $N_{max}$ replicates is compared to $\alpha$.

### Modification of CARAT

In order to assess the variance under the null of no association, CARAT uses a retrospective model for the tested marker $\mathbf{G}$ (see equation (14) in [4]). When we include the top PCs as covariates in our analysis, the GRM $\tilde{\mathbf{\Phi}}$ only reflects the leftover genetic correlation that is not captured by the top PCs. To incorporate all the genetic correlation, we consider the following retrospective model,

$$\mathbb{E}(\mathbf{G}|\mathbf{Y},\mathbf{Z}) = \mathbf{Z}\boldsymbol{\alpha} \text{ and } \mathrm{Var}(\mathbf{G}|\mathbf{Y},\mathbf{Z}) = \sigma_G^2\tilde{\mathbf{\Phi}}, \tag{5}$$

where $\mathbf{Z}$ is a $n \times (D+1)$ matrix whose columns are the top $D$ PCs along with an intercept, and $\boldsymbol{\alpha}$ is an unknown vector for their effects. The model in (5) allows for different MAFs conditional on each subject's genetic ancestry captured in the top PCs. Hence, we modify CARAT and use $\widetilde{\sigma}_G^2 = \mathbf{G}^T\mathbf{P}\mathbf{G}/(n-D-1)$, with $\mathbf{P} = \tilde{\mathbf{\Phi}}^{-1} - \tilde{\mathbf{\Phi}}^{-1}\mathbf{Z}(\mathbf{Z}^T\tilde{\mathbf{\Phi}}^{-1}\mathbf{Z})^{-1}\mathbf{Z}^T\tilde{\mathbf{\Phi}}^{-1}$, as an estimate of the variance parameter.

### P-values for IE and ED loci from other permutation-based methods

As described in Results, for IE, we detect a previously-identified [5] 12Mb region on chromosome 4 (position 7.5-19.3 Mb) that reaches the BRASS genome-wide significance threshold, with nominal p-value $2.1\times10^{-8}$ corresponding to a genomewide p-value of 8e-4 obtained from BRASS. The genomewide p-values for this variant by the other 6 methods, Naive, MVNpermute, LogMM-PQL, BRASS$_{mod}$, Naive$_{mod}$, and MVNpermute$_{mod}$, are, respectively, 4e-4, 7e-4, 2e-4, 7e-4, 1e-4, and 6e-4, based on 10,000 replicates of each. For ED, we detect 3 variants. The first is SNP rs9000666, which has nominal p-value 1e-7 and BRASS p-value 1e-2. The genomewide p-values for this variant by the other 6 methods, Naive, MVNpermute, LogMM-PQL, BRASS$_{mod}$, Naive$_{mod}$, and MVNpermute$_{mod}$, are, respectively, 4e-3, 6e-3, 3e-3, 7e-3, 4e-3, and 6e-3. The second is SNP rs21895578, which has nominal p-value 1.5e-7 and BRASS p-value 1.4e-2. The genomewide p-values for this variant by the other 6 methods, Naive, MVNpermute, LogMM-PQL, BRASS$_{mod}$, Naive$_{mod}$, and MVNpermute$_{mod}$, are, respectively, 5e-3, 9e-3, 4e-3, 1e-2, 6e-3, and 8e-3. The third is SNP rs23910667, which has nominal p-value 3.6e-7 and BRASS p-value .033. The genomewide p-values for this variant by the other 6 methods, Naive, MVNpermute, LogMM-PQL, BRASS$_{mod}$, Naive$_{mod}$, and MVNpermute$_{mod}$, are, respectively, .013, .021, .011, .025, .014, and .020.

### References

1. Balding DJ, Nichols RA. A method for quantifying differentiation between populations at multi-allelic loci and its implications for investigating identity and paternity. Genetica. 1995;96(1-2):3–12. doi:10.1007/BF01441146.

2. Marchini J, Cardon LR, Phillips MS, Donnelly P. The effects of human population structure on large genetic association studies. Nature Genetics. 2004;36(5):512–517. doi:10.1038/ng1337.

3. Conomos MP, Reiner AP, Weir BS, Thornton TA. Model-free Estimation of Recent Genetic Relatedness. American Journal of Human Genetics. 2016;98(1):127–148. doi:10.1016/j.ajhg.2015.11.022.

4. Jiang D, Zhong S, McPeek MS. Retrospective Binary-Trait Association Test Elucidates Genetic Architecture of Crohn Disease. American Journal of Human Genetics. 2016;98(2):243–255. doi:10.1016/j.ajhg.2015.12.012.

5. Hayward JJ, Castelhano MG, Oliveira KC, Corey E, Balkman C, Baxter TL, et al. Complex disease and phenotype mapping in the domestic dog. Nature Communications. 2016;7. doi:10.1038/ncomms10460.
