## Supplementary material for "BRASS: permutation methods for binary traits in genetic association studies with structured samples": S1 Table

**S1 Table. Differences between the seven resampling methods compared in the simulation studies.**

| Method | Feature: Binary trait replicate? | Trait variance is function of mean? | Resampling accounts for correlation in $\mathbf{Y}$? | Resampling accounts for correlation due to parameter estimation? |
|---|---|---|---|---|
| BRASS | – | $\surd$ | $\surd$ | $\surd$ |
| $\text{BRASS}_{mod}$ | $\surd$ | $\surd$ | $\surd$ | $\surd$ |
| LogMM-PQL | $\surd$ | $\surd$ | $\surd$ | – |
| MVNpermute | – | – | $\surd$ | $\surd$ |
| $\text{MVNpermute}_{mod}$ | $\surd$ | – | $\surd$ | $\surd$ |
| Naive | – | – | – | – |
| $\text{Naive}_{mod}$ | $\surd$ | – | – | – |

Check marks ($\surd$) indicate that the method incorporates the specified feature.
