## Supplementary figures and images for "BRASS: permutation methods for binary traits in genetic association studies with structured samples"

### S1 Fig

**S1 Fig. Three-generation pedigree used in the simulation studies.**

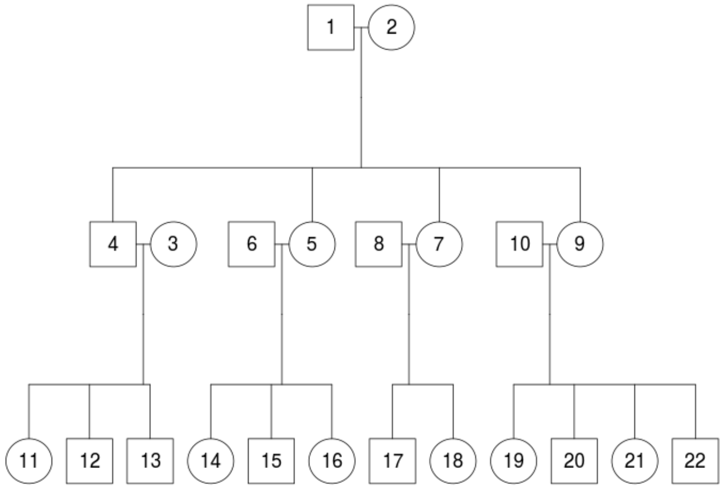

### S2 Fig

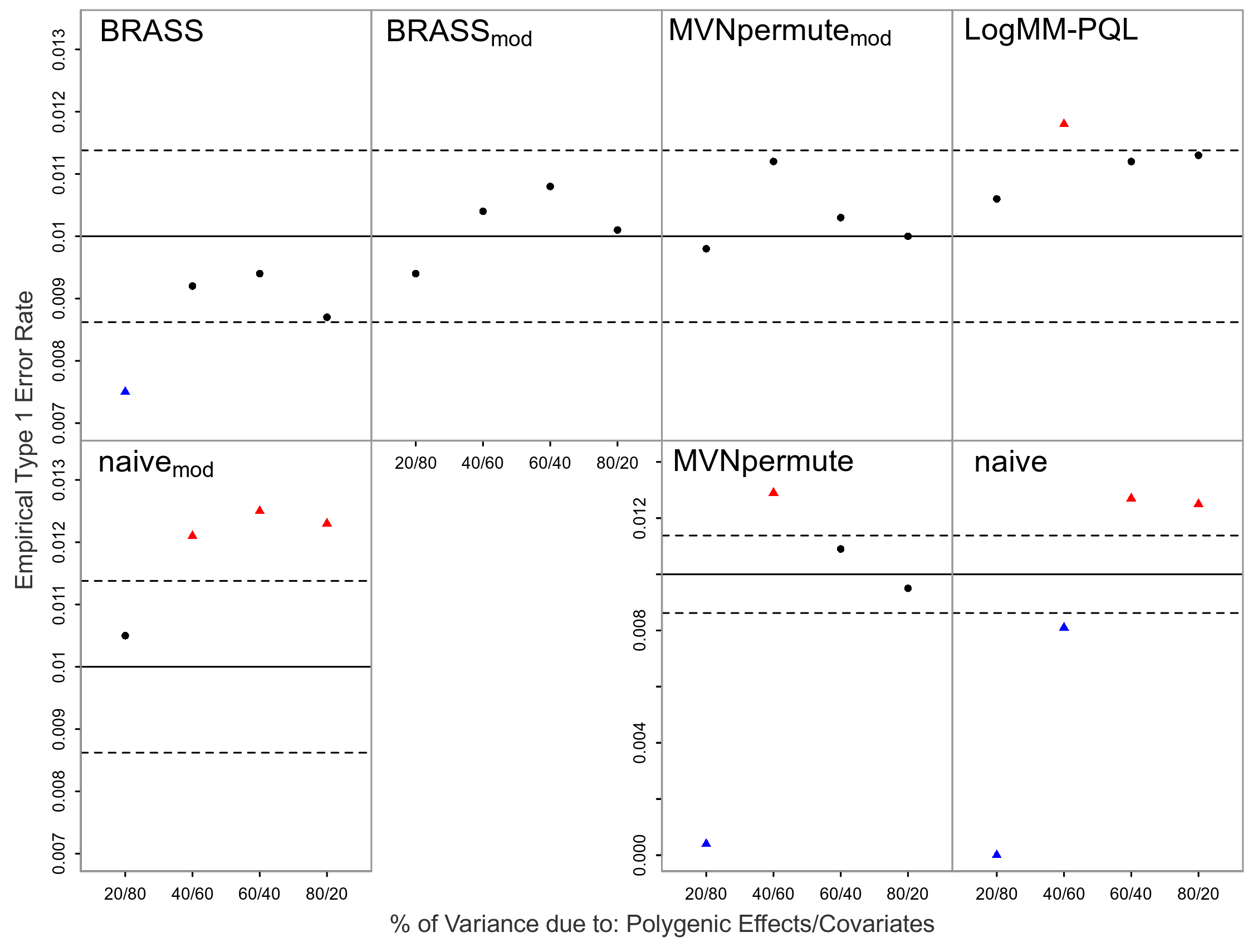
